# Effects of floral sex allocation and phenology on the within-flower selfing rate and female reproductive success in *Pulsatilla alpina*, a perennial herb with strong inbreeding depression

**DOI:** 10.1101/2024.05.12.593796

**Authors:** Kai-Hsiu Chen, John R. Pannell

## Abstract

1. Within-flower self-pollination should be the major source of self-fertilization in mixed-mating species presenting single or few flowers simultaneously. It is also an often unmeasured source of selfing in species with many flowers open simultaneously. In general, the rate of within-flower selfing is expected to depend on the number of pistils and stamens, the timing of flowering, and subsidiary floral traits in self-compatible species in which pistil and stamen numbers vary. Accordingly, the intensity and direction of selection on these traits should depend on the level of inbreeding depression.
2. Here, we measured the dependence of the within-flower selfing rate on floral sex allocation, phenology, petal length, and floral stalk height in a population of the perennial herb *Pulsatilla alpina* (Ranunculaceae) in which most individuals had single flowers. We estimated inbreeding depression in the population by comparing inbreeding coefficients between parents and seed progeny using microsatellite markers. We then estimated selection on the measured traits via female reproductive success at the flower level and compared our estimates with a hypothetical scenario in which inbreeding depression was assumed to be absent.
3. We estimated inbreeding depression to be 0.93. The within-flower selfing rate varied widely among flowers and depended positively on stamen number and negatively on pistil number and flowering date, supporting the predictions of a mass-action model. The dependence of the selfing rate on the measured floral traits consistently predicted (non-linear) patterns of selection under high inbreeding depression that were distinct from those under a hypothetical scenario of no inbreeding depression.
4. *Synthesis:* While previous research has emphasized the importance of mass-action mating on selfing among flowers of plants with large floral displays, our results demonstrate its importance for selfing within individual flowers. They also demonstrate the importance of accounting for both the selfing rate and inbreeding depression when inferring selection on floral and other traits via female fitness.

## Introduction

The rate of self-fertilization in self-compatible plants and the level of inbreeding depression experienced by progeny derived from self-fertilization (hereafter ‘selfed progeny’) are major determinants of female reproductive success. Traits that affect the selfing rate are thus expected to be under strong selection. One such trait is the spatial separation between the sexual parts (herkogamy) (Webb & Lloyd, 1986; Barrett et al., 2003), with a negative correlation between the degree of herkogamy and the selfing rate having been found in at least 20 species (reviewed in Opedal, 2018). Another such trait is the relative timing of stigma receptivity and pollen release from anthers in the same flower (dichogamy) (Lloyd & Webb, 1986), e.g., reduced temporal separation between the two sex functions was found to be positively correlated with the selfing rate in *Campanula americana* (Koski et al., 2018) and *Gilia achilleifolia* (Schoen, 1982). Almost all evidence for the effects of herkogamy and dichogamy is drawn from comparisons among populations that vary in their mean levels of spatial or temporal separation between the sexes (Koski et al., 2018; Opedal, 2018; Schoen, 1982), and there is a surprising dearth of evidence for variation in the selfing rate as a function of variation in these factors among individuals within populations, where ultimately natural selection will be strongest.

In addition to herkogamy and dichogamy, a flower’s sex allocation, i.e., the absolute and/or relative numbers of pistils and stamens, should also be expected to affect the rate of within-flower selfing (including both autonomous and pollinator-facilitated components of selfing). Although species in most of the monocots and core eudicots tend to have fixed numbers of pistils and stamens, many species from basal clades (e.g., basal angiosperms and Magnoliids) do have variable numbers of floral parts (Ronse de Craene et al., 2003), and this variation could affect the selfing rate in self-compatible species (Allen & Hiscock, 2008; Barrett, 2013; Igic et at., 2006). Furthermore, many species present single or few flowers daily or throughout a flowering season, and within-flower selfing should be the dominant source of selfing in these species (e.g., Karron et al., 2004; Méndez & Traveset, 2003; Sun et al., 2018). In these species, we should expect flowers with more stamens per pistil to have a higher selfing rate, just as selfing due to pollen transfer among flowers of the same individuals (geitonogamy) has been shown to depend positively on the number of flowers in the floral display (Harder & Barrett, 1995; Karron et al., 2004; Williams, 2007). The idea that sex allocation should affect the selfing rate is central to a number of models of mating-system or sex-allocation evolution (Brunet & Charlesworth, 1995; de Jong et al., 1999; Gregorius et al., 1987; Holsinger, 1991; Lloyd, 1992; Sakai, 1995). However, although there has been substantial empirical work on the effects of geitonogamy and sex allocation on the selfing rate at the individual level (Harder & Barrett, 1995; Karron et al., 2004; Williams, 2007), we are not aware of any study that has quantified the relationship between within-flower sex allocation and the selfing rate, despite its likely importance for the evolution of floral morphology and sex allocation.

The within-flower selfing rate should depend not only on the numbers of stamens and pistils in flowers of species that vary for these traits, but also on the timing of their presentation to pollinators and of the presentation of the other flowers in a population. If mating follows a ‘mass-action’ process, in which the mating system depends on the relative availability of self versus outcross mates at a given point in time (Gregorius et al., 1987; Holsinger, 1991), we may expect a decrease in the selfing rate in flowers that produce fewer stamens and that are presented to pollinators when the density of other flowering individuals is high. For example, in the semi-desert plant *Incarvillea sinensis* and the alpine plant *Phyllodoce aleutica*, an elevated selfing rate was found in the late and early seasons, respectively, when pollinator visitation rates were low (Kameyama & Kudo, 2009; Yin et al., 2016). However, we are largely ignorant of how temporal variation in plant mating systems relates to quantitative variation in the timing of male and female functions over the course of a flowering season.

Natural selection should act on floral morphology and on their sex allocation and phenology as a function of the selfing rate and the level of inbreeding depression suffered by selfed progeny. Numerous studies have considered the effect of different floral traits on the female component of fitness (reviewed in Ashman & Morgan, 2004; Munguía-Rosas et al., 2011; Caruso et al., 2019), but these studies are typically based on numbers of seeds produced, without accounting for the possibility that many seeds might be self-fertilized and could suffer inbreeding depression (Lloyd, 1992). Nor have any of these studies considered the effects of within-flower selfing as opposed to selfing that includes geitonogamy. Although within-flower selfing and inter-flower selfing are genetically identical, they have different implications for the evolution of floral traits. For instance, a reduction in floral display by presenting fewer flowers in an inflorescence simultaneously might be an effective mechanism to avoid geitonogamous selfing (Harder & Barrett, 1995; Karron et al., 2004; Williams, 2007), while such a mechanism would have very little impact on within-flower selfing. If we are to gain a balanced understanding of how selection shapes the timing and amount of allocation to male and female functions within flowers, we need to consider how these traits affect not only the number of seeds produced but also their quality (as well as the male components of reproductive success). To our knowledge, nearly all the studies of phenotypic selection have not accounted for the potential consequence of inbreeding, and those few that did were conducted in artificial experimental arrays with an unknown level of inbreeding depression (e.g., Briscoe Runquist et al., 2017; Hou et al., 2024). How mating-system variation and the consequence of inbreeding depression alters selection on different traits in wild populations thus remains an open question.

Here, we ask how the amount and timing of sex allocation at the flower level affects within-flower selfing and, consequently, the female component of seasonal reproductive success. We estimated the selfing rate of progeny within flowers and the level of inbreeding depression suffered by selfed progeny in a population of the insect-pollinated and self-compatible species *Pulsatilla alpina* to determine how the selfing rate and female reproductive success depends on floral sex allocation, phenology, and subsidiary floral traits. Because the study population comprised mainly individuals with only one flower, we could assign paternity at the flower level and thus relate contributions to fitness to the traits of individual flowers. Although flowers vary substantially in their pistil and stamen number, we further enhanced this variation by removing stamens from a random subset of flowers in the population, a treatment that also generated purely female flowers that do not otherwise occur in *P. alpina.* We estimated the female component of seasonal reproductive success by counting seeds produced by individual flowers through both outcrossing and within-flower selfing, accounting for the negative effects on fitness of inbreeding depression. We then quantified phenotypic selection on the traits under two scenarios of inbreeding depression. We specifically addressed the following questions. 1) How does the within-flower selfing rate depend on the number of pistils and stamens within flowers, the timing of flowering, and two subsidiary floral traits? 2) What is the level of inbreeding depression? 3) How does the inbreeding depression affect the pattern of selection on the traits as a result of the dependency of within-flower selfing on those traits?

## Materials and Methods

### Study species and study sites

*Pulsatilla alpina* (L.) Delarbre (Ranunculaceae) is a perennial herb that grows in sub-alpine to alpine habitats in central Europe (Lauber et al., 2018). It persists over winter as an underground stem (rhizome) and emerges soon after the snowmelt, from early May to July, in the form of several vegetative and/or reproductive shoots. Depending on their size and age, individuals produce a mixture of up to 20 male and bisexual flowers (on average 2.05 ± 1.76 flowers based on a sampling of 1131 individuals across 12 populations in 2022; Figure S1), each on its own reproductive shoot and each with around six showy white tepals (the petals and sepals are not distinguishable). Male flowers bear only stamens, whereas bisexual flowers bear stamens and one to a few hundred uni-ovulate pistils (Figure 1). Bisexual and male flowers bear a similar number of stamens (Chen & Pannell, 2023a). Bisexual flowers are strongly protogynous, with pistils receptive before pollen is dispersed from stamens. The species is self-compatible and likely capable of self-fertilizing autonomously without pollinators (Figure S2). Individuals have size-dependent sex allocation, with larger plants allocating absolutely and proportionally more resources to their female function (Chen & Pannell, 2023a). Small individuals often produce only a single male flower and thus function as pure males in the respective flowering season (Chen & Pannell, 2023a), leading to ‘gender diphasy’ (Schlessman, 1988). Both male and bisexual flowers are visited predominantly by dipteran insects, including houseflies and syrphid flies (Chen & Pannell, 2022). Ripe fruits (technically ‘achenes’) with elongated pappus hair are dispersed from the extended floral stalks by wind in early autumn (Chen & Pannell, 2022; Vittoz & Engler, 2007). After seed dispersal, above-ground vegetative parts senesce.

**Figure 1.**
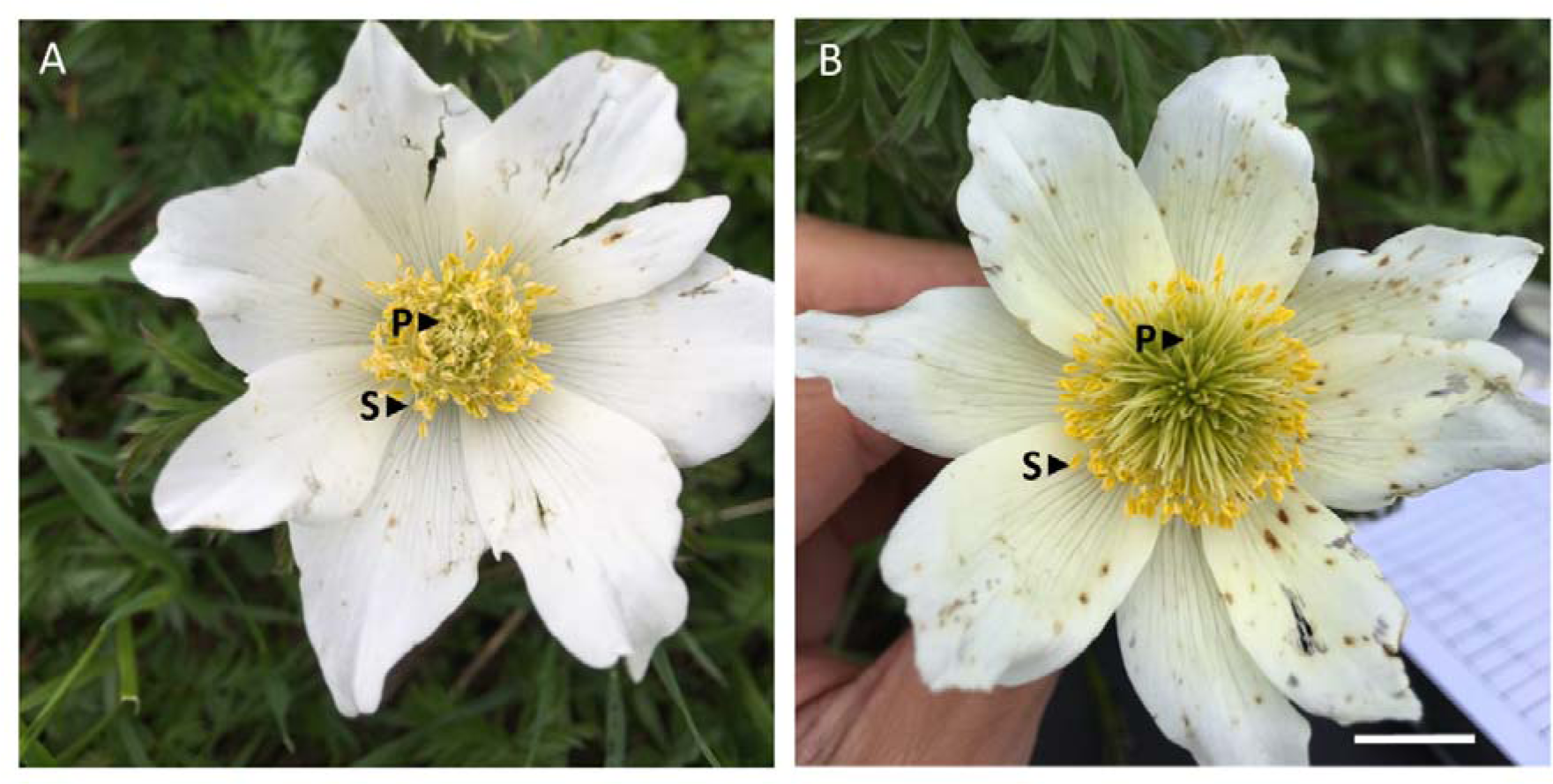
Photographs of hermaphroditic flowers of *P. alpina* with a low (A) and high (B) number of pistils. The pistils (P) are in the inner whorls, surrounded by numerous stamens (S). The white bar represents 2 centimeters.

In the flowering season of 2022, we studied a single population of *P. alpina*, located at Solalex in the pre-Alps of Vaud canton, Switzerland (‘Population S1+’; latitude: 46°17′42″N, longitude: 7°09′09″E; elevation: 1758 a.s.l.). The population was located on an open slope of sub-alpine grassland and covered an area of about 20 m x 20 m, comprising about 150 mainly small and probably young individuals (following recent establishment after avalanche disturbances and/or herbivory by cattle). Most individuals in the population produced only a single flower in 2022 (and consecutive years; unpublished data), providing an unusual opportunity to study the within-flower selfing rate without having to account for inter-floral selfing. We considered within-flower selfing as a joint consequence of autonomous selfing and pollinator-facilitated selfing, because the two modes of selfing occur simultaneously during flowering. We set up a 10 m x 15 m temporally fenced plot within the population, enclosing 135 flowering individuals, and removed all the floral buds outside the plot (i.e., around 10) prior to the onset of flowering to prevent nearby individuals from siring progeny in the plot.

### Measurements of floral traits

We measured two sex allocation traits (stamen number and pistil number) and three display traits (flowering date, tepal length, and floral stalk height) for all flowers in our sample (*N* = 135). We visited the population every three to four days throughout the flowering season, from late May to late June 2022, noting the number of flowers and stalks, and the height of the tallest foliar stalk of all flowering individuals. Flowers were individually labelled with a paper tag. We photographed each sampled flower at the end of the female stage (see Chen and Pannell, 2023b for a detailed description of the categories), and later counted stamen number on the basis of the photographs (see next section below). The flowering date for each flower was set as the date of its opening. Tepal length and floral stalk height were measured at the end of the male stage (following the method used by Chen and Pannell, 2022). Around three weeks after the end of the flowering season, all flowers with developing achenes were enclosed in a paper bag until the end of the growing season (early August), at which point all their achenes were collected. The pistil number was calculated as the total number of achenes collected because all the pistils elongated and remained on the floral stalk, whether pollinated or not (see details below).

### Manipulation of stamen number

To increase variation in male allocation among bisexual flowers, we allocated a subsample of individuals to one of two stamen-removal treatments (c.f. Emms, 1993; e.g., Yund, 1998; Johnson & Yund, 2009; Aljiboury & Friedman, 2022). Specifically, we randomly selected about a quarter of the bisexual flowers and removed either 100% or 50% of their stamens before any of them had dehisced (treatments SR_100_ or SR_50_, respectively). These treatments had no effect on the flowering duration of the affected flowers (Chen & Pannell, 2023b). We counted the actual number of stamens for 15 flowers in vivo (*N*_actual_) as well as with the aid of photographs (*N*_photo_), and we used the regression *N*_actual_ = 1.66 x *N*_photo_ + 27.2 (*r*^2^ = 0.654) (Chen & Pannell, 2023a) to calibrate our stamen counts based on photographs for flowers of intact and SR_50_ treatments. We used the number of stamens as a proxy for male allocation by assuming that each stamen produces the same amount of pollen (but see Murakami et al., 2022; Spalik & Woodell, 1994).

### Estimation of the selfing rate

We estimated the selfing rate *S_i_* of each bisexual flower *i* based on variation at ten microsatellite loci, as described in Chen and Pannell (2023b). Briefly, we extracted DNA from 1,054 mature seeds (an average of 10.2 ± 1.3 seeds per family) and 135 parental individuals. A total of 892 seeds that could be genotyped for at least five loci were ultimately used for paternity analysis. Of these, we were able to assign the paternity of 854 seeds (96%) to a single most likely father under a relaxed confidence interval (80%). The within-flower selfing rate was calculated for each seed family by dividing the number of selfed seeds by the number of successfully genotyped seeds.

### Estimation of inbreeding depression

We used variation estimated at the same ten microsatellite loci to calculate inbreeding coefficients for adults and seed progeny (*F*_p_ and *F*_o_, respectively), using the R package ‘*hierfstat*’ (Goudet, 2005), and estimated inbreeding depression by comparing the two coefficients, following Ritland (1990). This approach assumes that a reduction in *F* from the progeny to parental generations is due to differential mortality of selfed progenies as a result of inbreeding depression. The population-level selfing rate, *S*, was estimated by dividing the sum of the inferred number of selfed seeds of each seed family by the total number of mature seeds of the population. Inbreeding depression was then estimated as below:

*d* = 1 - [2(1- *S*)*Fp⁄S*(1+ *F_o_* - 2*Fp*)] (Ritland, 1990).

### Estimation of female reproductive success

We estimated female reproductive success as the number of gene copies potentially transmitted to the next generation through seeds. We collected achenes of all bisexual flowers that had not been aborted by the end of the flowering season and sorted them into unfertilized, predated, and mature categories based on their morphology, following Chen and Pannell (2022); note that each pistil contains only one ovule and that the sum over all three categories represents a flower’s total pistil number. We then calculated female reproductive success based on our estimated value of inbreeding depression, *d*, as well as assuming *d* = 0; the comparison of reproductive success estimated under these two scenarios allowed us to infer the extent to which phenotypic selection depends on inbreeding depression. We calculated female reproductive success for our estimate of *d* as the number of mature outcrossed seeds plus (1 – *d*) times the number of mature seeds produced by selfing. When *d* = 0, female reproductive success equals the number of mature seeds, as estimated in most empirical studies that have measured phenotypic selection via female reproductive success without considering the mating system. When *d* > 0, the reduced fitness of selfed progeny reduces the fitness of both the female and the male components of fitness via selfing (e.g., Briscoe Runquist et al., 2017; Hou et al., 2024). Many theoretical studies attribute the two-fold transmission advantage of selfing solely to female reproductive success, i.e., doubling the female contribution of selfed seeds (e.g., Charlesworth & Charlesworth, 1978; Lloyd, 1987). We decomposed fitness components in this way, too (see Figure S3 for these results), but because the inferred patterns of phenotypic selection were almost identical, we present results in the main text based on assigning fitness under selfing to both female and male reproductive success.

### Statistical analysis

We conducted all analyses within the *R* statistical framework v4.0.3 (*R* Core Team, 2021). We evaluated the fit of the linear models with the *R* package *DHARMa* (Hartig, 2019). We used a generalized linear mixed model (*glmer* function in the *R* package *lmer*; Bates et al. 2015) to evaluate the dependence of the within-flower selfing rate on five floral traits for individuals producing only one bisexual flower with at least six successfully genotyped seeds. We set the within-flower selfing rate as a binomial response variable. We standardized pistil number, stamen number, flowering date, stalk height, and tepal length to a mean of zero and a standard deviation of one and set each of the standardized traits as explanatory variables. We set the identity of each flower as a random variable to account for the fact that the genotyped seeds from the same flower were not independent of each other in terms of the floral traits shared. The *P*-values were calculated using likelihood ratio tests. The variance explained by the model was checked using the *R* package *MuMIn* (Barton, 2009).

We used two multivariate regression models to evaluate how the contrasting inbreeding depression scenarios affect phenotypic selection on five floral traits via the two estimates of female reproductive success. First, we used a linear mixed model with relative female reproductive success as a response variable, with a Gaussian error distribution. Second, we used a generalized linear mixed model with mature seed number as a response variable, with a Poisson distribution. Both models represent two commonly used approaches to quantify the dependence of components of reproductive success on phenotypic traits (Lande & Arnold, 1983; Morrissey & Goudie, 2022); the latter has been widely applied in recent studies in plants to study phenotypic selection (e.g., Fournier-Level et al. 2022; Marrot et al. 2022; Castellanos et al. 2023). Rather than comparing the performance of the two models, our aim here was to evaluate two perspectives for evidence of interactive effects between inbreeding depression scenarios and regression coefficients (see Morrissey and Goudie, 2022, for the comparison between the two approaches).

For the two regression models, we included all individuals with a single bisexual flower in the population and a complete set of floral trait measurements. In the linear mixed model, we calculated the relative seasonal female reproductive success by dividing the number of mature seeds of each individual by the mean number of mature seeds across the individuals considered, as defined by Lande and Arnold (1983), with the five floral traits standardized, as described above. To estimate linear and quadratic selection coefficients, which represent direct selection on the trait (i.e., the partial effect), we set linear and quadratic terms for the five traits as fixed effects and considered the interactions with the degree of inbreeding depression (i.e., *d* = 0 or 0.93) in the model (Lande & Arnold, 1983; Matsumura et al., 2012). We used the *emtrends* function in the *R* package *emmeans* to extract from the model the regression coefficients and standard errors of the traits under the two inbreeding depression scenarios (Lenth, 2020). A significant linear coefficient indicates a directional selection on the trait, while a significant quadratic coefficient in conjunction with a local maximum and minimum indicates stabilizing or disruptive selection, respectively. We used likelihood ratio tests to extract the *P*-values for the interaction terms. A significant interaction between a floral trait and inbreeding depression scenarios indicates a difference in the pattern of selection. We set individual identity as a random effect to account for the fact that the estimates of relative reproductive success of the same individual under the two scenarios are not independent.

In the generalized linear mixed model, the mature seed number estimated under the two inbreeding depression scenarios was used as the response variable, with the same model structure of fixed and random effects described for the previous model. The regression coefficients and standard errors for each trait were extracted from the model using the *emtrends* function in the *R* package *emmeans* under the two inbreeding depression scenarios (Lenth, 2020). We used likelihood ratio tests to calculate the *P*-values for the interaction terms.

## Results

### Within-flower selfing and floral traits

The within-flower selfing rate varied substantially among single-flowered individuals, with a mean of 0.45 and a standard deviation of 0.39 (*N* = 621 seeds from 53 individuals; see Figure S4 for the distribution). The within-flower selfing rate increased with the flower’s number of stamens (*P* < 0.001; Figure 2A), decreased with its pistil number (*P* < 0.05; Figure 2B) and flowering date (*P* < 0.05; Figure 2C), and was independent of its tepal length and stalk height (*P* > 0.05; Table S2). Means and standard deviations of the five floral traits as well as the pairwise correlations between traits for the single-flowered individuals sampled are provided in Table S1 and Figure S5. Our model explained 71% of the variance in the observed selfing rate (conditional *R^2^*).

**Figure 2.**
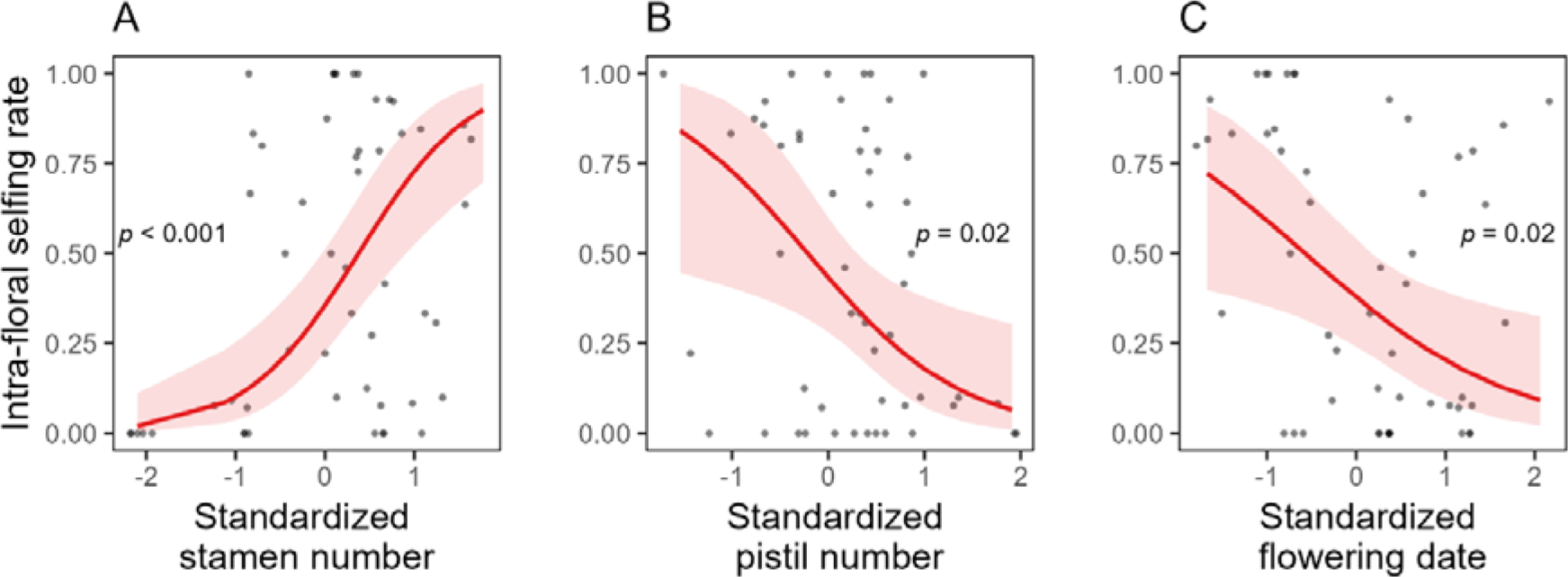
Plots showing the partial effect of stamen number (A), pistil number (B), and flowering date (C) on the within-flower selfing rate of single-flowered hermaphroditic individuals of *P. alpina,* estimated by a multivariate *glmer* model (*N* = 621 genotyped seeds from 53 individuals). The data points were jittered to avoid overlapping. The shaded ribbon indicates the 95% confidence interval of the regression curves.

### Estimate of inbreeding depression

The inbreeding coefficient of the parents, *F*_p_ (*N* = 135 flowering individuals), and offspring, *F*_o_ (*N* = 1054 seeds), were 0.028 and 0.221, respectively, implying an inbreeding depression of 0.93 (Ritland, 1990).

### Phenotypic selection coefficients under two scenarios of inbreeding depression

Table 1 presents the direction and intensity of linear and quadratic coefficients on five floral traits for the estimated value of inbreeding depression in the population (0.93) and the hypothetical contrasting value of 0. Using relative female reproductive success as the proxy for fitness, interaction terms between the degree of inbreeding depression and selection coefficients were significant for pistil number (both linear and quadratic gradients), stamen number (quadratic gradient), and flowering date (quadratic gradient; marginally significant) (Table 1; Figure 3A, B, and C). Using mature seed number as the proxy for fitness, interaction terms were significant for both linear and quadratic coefficients in pistil number, stamen number, flowering date, and tepal length (Table 1; Figure 3F, G, H, and I). For both fitness proxies, taking inbreeding depression into account increased the quadratic coefficients on pistil number, revealed mostly negative directional selection on stamen number, and shifted the direction of the quadratic coefficients on flowering date (Table 1; Figure 3F, G, H, and I). See Chen and Pannell (2023b, 2024) for how the traits affected different components of male siring success.

**Figure 3.**
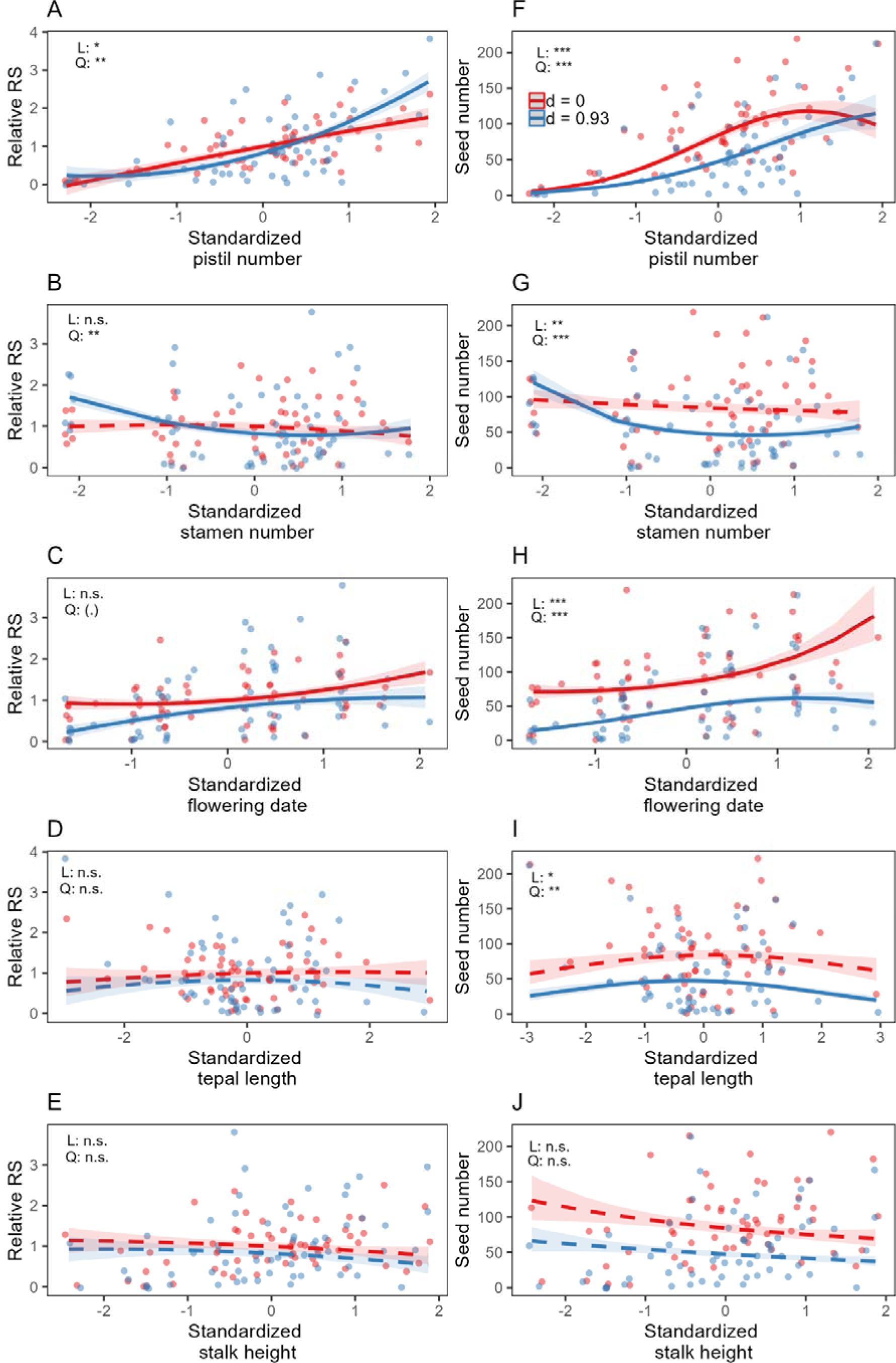
Plot showing the dependence of female fitness on pistil number (A, F), stamen number (B, G), flowering date (C, H), tepal length (D, I), and stalk height (E, J) of single-flowered hermaphroditic individuals of *P. alpina*, using relative reproductive success (relative RS, left-hand panel) and mature seed number (right-hand panel) as fitness proxies (*N* = 60 individuals). Each individual is represented by two points for scenarios of an inbreeding depression of zero (red) and 0.93 (blue). The data points were jittered to avoid overlapping. The shaded ribbons indicate the standard error of the regression curves. Dashed regression curves indicate non-significant linear and quadratic coefficients. The *P* values of interaction terms of inbreeding depression scenarios with linear (L) and quadratic (Q) coefficients are denoted by asteroids. See Table 1 for details of the values and statistics of the linear and quadratic coefficients.

**Table 1.** Linear and quadratic coefficients via the female function on five floral traits on single-flowered hermaphroditic individuals of *P. alpina* under two scenarios of inbreeding depression in models using relative reproductive success and mature seed number as the proxy for female fitness (*N* = 60). Standard errors are given in parentheses. A significant interaction indicates that the selection coefficients under the two scenarios are statistically different. Notes: (●) *P* < 0.1, ∗ *P* < 0.05, ∗∗ *P* < 0.01, ∗∗∗ *P* < 0.001

| Model | Coefficient | Floral trait | $d = 0$ | $d = 0.93$ | Interaction |
| --- | --- | --- | --- | --- | --- |
| A. Relative RS | Linear term | Stamen number | -0.07 (0.1) | -0.15 (0.1) |  |
|  |  | Pistil number | 0.42 (0.11) *** | 0.65 (0.11) *** | * |
|  |  | Flowering date | 0.17 (0.09) (●) | 0.25 (0.09) ** |  |
|  |  | Tepal length | 0.04 (0.08) | -0.003 (0.08) |  |
|  |  | Stalk height | -0.09 (0.1) | -0.1 (0.1) |  |
|  | Quadratic term | Stamen number | -0.03 (0.07) | 0.13 (0.07) (●) | ** |
|  |  | Pistil number | -0.01 (0.07) | 0.17 (0.07) * | ** |
|  |  | Flowering date | 0.08 (0.09) | -0.06 (0.09) | (●) |
|  |  | Tepal length | -0.01 (0.05) | -0.03 (0.05) |  |
|  |  | Stalk height | -0.01 (0.07) | -0.02 (0.07) |  |
| B. Seed number | Linear term | Stamen number | -0.05 (0.08) | -0.14 (0.08) (●) | ** |
|  |  | Pistil number | 0.6 (0.09) *** | 0.77 (0.09) *** | *** |
|  |  | Flowering date | 0.22 (0.07) ** | 0.43 (0.07) *** | *** |
|  |  | Tepal length | 0.01 (0.07) | -0.05 (0.07) | * |
|  |  | Stalk height | -0.13 (0.08) | -0.13 (0.08) |  |
|  | Quadratic term | Stamen number | 0.005 (0.06) | 0.14 (0.06) * | *** |
|  |  | Pistil number | -0.27 (0.06)<br>** | -0.16 (0.06)<br>*** | *** |
|  |  | Flowering date | 0.07 (0.07) | -0.17 (0.08) * | *** |
|  |  | Tepal length | -0.04 (0.04) | -0.09 (0.04) * | ** |
|  |  | Stalk height | 0.01 (0.06) | 0.002 (0.06) |  |

## Discussion

Our results confirm that *Pulsatilla alpina* is self-compatible and has a mixed mating system, with an average within-flower selfing rate in the study population close to 0.45. This estimate is comparable with that recorded for other self-compatible perennial herbs pollinated by insects (references in Whitehead et al., 2018), particularly plants with open flowers and visited by generalist pollinators such as flies (as is the case for *P. alpina*) (Shirk & Hamrick, 2014; Tamaki et al., 2019; Wirth et al., 2010). Although individuals in many *P. alpina* populations produce several flowers simultaneously (Figure S1), so that geitonogamy may contribute to the selfing rate, there was in fact no difference in our estimate of the selfing rate between individuals with one flower and those producing several (Chen & Pannell, 2023b). Geitonogamy thus appears to contribute little to the overall selfing rate of the study population. The flowers of *P. alpina* are actinomorphic and contain several whorls of pistils surrounded by several whorls of stamens, with anthers and stigmas positioned at the same level, i.e., flowers lack any substantial herkogamy (or ‘unordered’ herkogamy *sensus* Webb and Lloyd, 1986). Although flowers are strongly protogynous, with acropetal maturation of floral parts, it is clear that the temporal separation of the sex functions within flowers does not completely prevent selfing (Figure S2).

We found that the within-flower selfing rate of *P. alpina* increased with allocation to male function and decreased with allocation to female function. It seems likely that stigmas of the outer whorl of pistils are still receptive when they receive pollen from the anthers of the inner whorl of stamens, and that the production of more pistils and fewer stamens within a flower reduces the proportion of stigmas close to stamens, thereby increasing the average degree of herkogamy within flowers. There is now substantial evidence from studies of species with a fixed number of stamens and pistils that the selfing rate is a negative function of the degree of herkogamy, i.e., anther-stigma separation (e.g., Brunet & Eckert, 1998; Takebayashi et al., 2005; Medrano et al., 2012; reviewed in Opedal, 2018). Our results indicate that, in species with a variable number of floral parts, i.e., in most basal angiosperms and basal eudicots that have multiple whorls of sexual organs (Kim et al., 2005; Méndez & Traveset, 2003; Ronse de Craene et al., 2003), herkogamy may also depend indirectly on the relative numbers of pistils and stamens. This indirect effect of sex allocation on herkogamy has, to our knowledge, not previously been recorded.

Another key result of our study is that the within-flower selfing rate of *P. alpina* decreased with time over the flowering season. This pattern is consistent with assumptions of mass-action models that assume that the selfing rate depends on the contribution to local pollen production of both the focal plant (which increases the selfing rate) and the pollen production of neighbouring plants (which dilutes the focal plant’s pollen and reduced the selfing rate). Mass action assumptions have been assumed in many models of plant mating and sexual systems (Brunet & Charlesworth, 1995; de Jong et al., 1999; Gregorius et al., 1987; Holsinger, 1991; Lloyd, 1992; Sakai, 1995). Our results indicate that such models might be a reasonable approximation of mating patterns for species beyond those pollinated by wind, at least those pollinated by generalist pollinators such as flies, as is the case for *P. alpina*. How the mating system depends on phenology at a fine scale has rarely been studied in alpine plants, despite the importance of phenology on their reproductive success (Inouye, 2020; Kudo, 2022; Straka & Starzomski, 2015), though a similar pattern of increased selfing rate in the early flowering season has been found in the bumblebee-pollinated alpine plant *Phyllodoce aleutica* (Kameyama & Kudo, 2009). In *P. aleutica*, plants at early snowmelt patches had a higher selfing rate compared to late snowmelt patches, likely due to a lack of pollinators early in the season (Kameyama & Kudo, 2009), as appears to be the case for *P. alpina* (although we did not quantify pollinator activity in this study).

Our comparison of the inbreeding coefficient among seeds (high) and among adults (low) points to very strong inbreeding depression in the study population, which we estimated at 0.93. High inbreeding depression is frequent in long-lived perennial plants (Angeloni et al., 2011; Goodwillie et al., 2005; Morgan, 2001; Scofield & Schultz, 2005), often expressed during the early stages of plant growth (Husband & Schemske, 1996). Edelfeldt et al. (2019) measured mortality for a related species of *Pulsatilla* and, similarly, inferred it to be highest at early stages of growth and then independent of age and size. The high level of inbreeding depression combined with substantial selfing observed in *P. alpina* implies strong ongoing selection for improved outcrossing mechanisms (Chen & Pannell, 2023b), as well as the existence of sexual conflict in the allocation of resources to male versus female functions. Indeed, the negative dependence of female reproductive success on male allocation, a sign of sexual interference, was revealed only when we accounted for the effect of inbreeding depression.

Sexual interference occurs when the presence of one sexual function decreases the reproductive success via the opposite sex (Barrett, 2002). In *P. alpina*, sexual interference is manifested through seed discounting due to a positive dependence of the within-flower selfing rate on stamen number. Previous empirical studies have inferred sexual interference in terms of a deleterious effect of male allocation on female reproductive success, but these have emphasized geitonogamous selfing as the key mechanism (e.g., Harder et al., 1995; Duffy & Johnson, 2014; Larue et al., 2022). Our results demonstrate that interference through sex allocation can also occur at the within-flower level. Such within-flower sex-allocation interference might be common in other mixed-mating species, especially those in which anther and pistil numbers vary.

The significant interactions of inbreeding scenarios with linear and quadratic selection on female allocation in *P. alpina* also have implications for the empirical estimation of female fitness gain curves (Campbell, 2000). Fitness gain curves trace the dependence of fitness gained through a given sex function on sex allocation and are thought to predict the evolutionary stable sex allocation strategy of the population (Charlesworth & Charlesworth, 1981; Charnov et al., 1976; Lloyd, 1984). We found that when taking high inbreeding depression into account, the female gain curve at the flower level became less saturating, because flowers that allocated more to their female function and less to their male function selfed less and thus suffered lower seed discounting (*sensu* Lloyd, 1992). Such a scenario leads to disruptive selection on sex allocation, favoring flowers with either no pistils or those with many pistils (Chen & Pannell, 2023b). This within-flower sexual interference in sex allocation in *P. alpina* provides a plausible explanation for the evolution of its strategy of andromonoecy, in which individuals produce either male or bisexual flowers (see the following empirical studies for tests of alternative but not mutually exclusive hypothesis: Granado-Yela et al, 2017; Kudo & Shibata, 2021; Miller & Diggle, 2003; Murakami et al., 2022; Tomaszewski et al., 2018).

We note that the selection gradient on the flowering date via female function was mostly directional, with flowers that allocated more resources to their female function selected to open later in the season, independent of the level of inbreeding depression in progeny. This finding differs from a review of data from 87 species that found that selection on flowers through their female function generally favored early flowering, especially in the temperate zone (Munguía-Rosas et al., 2011). The discrepancy is likely due to the fact that sex allocation in *P. alpina* is protogynous, with an emphasis on male function in early flowers and a tendency towards greater female allocation in flowers later in the season (Brunet and Charlesworth, 1995; Austen et al., 2015; Chen and Pannell, 2023a; Figure S5). In contrast, most of the species reviewed by Munguía-Rosas et al. (2011) are protandrous, for which we expect individuals to adopt a reciprocal strategy of phenology-dependent sex allocation, with flowers emphasizing their female function selected to open in the early season, and vice versa for flowers emphasizing their male function. Ultimately, we should expect selection on sex allocation and its timing within flowers to depend strongly on whether the species concerned is protandrous or protogynous.

Finally, our results prompt a degree of caution in the interpretation of selection gradient analyses using seed number as a fitness proxy for species that may have a mixed mating system and high inbreeding depression. Most empirical studies using seed number as the female fitness proxy have not accounted for inbreeding and its possible effects, despite the fact that many species have a mixed-mating system with high inbreeding depression (Caruso et al., 2019; Harder & Johnson, 2009; Munguía-Rosas et al., 2011). Our results demonstrate that ignoring the impact of inbreeding depression on components of fitness may lead to erroneous inferences, particularly on the intensity and direction of non-linear selection. This cautionary note should of course also apply to selection gradient analysis on species in which selfing results from geitonogamy (e.g., Van Kleunen & Ritland, 2004; Briscoe Runquist et al., 2017; Hou et al., 2024).

## Notes

### Competing Interest Statement

The authors have declared no competing interest.

