## Supplementary material for "Effects of floral sex allocation and phenology on the within-flower selfing rate and female reproductive success in *Pulsatilla alpina*, a perennial herb with strong inbreeding depression"

**Supplementary Tables and Figures**

|  | Mean (SD) |
| --- | --- |
| Floral traits |  |
| Stamen number (before manipulation) | 179.7 (35.9) |
| Stamen number (after manipulation) | 146.2 (69.5) |
| Pistil number | 215.6 (94.1) |
| Flowering date (Julian day) | 159.7 (7) |
| Tepal length (cm) | 2.81 (0.27) |
| Stalk height (cm) | 38.1 (7.5) |
| Female reproductive success |  |
| Mature seed number (*d* = 0) | 89.3 (52.6) |
| Mature seed number (*d* = 0.93) | 55.9 (49.6) |

**Table S1.** Mean and standard deviation (SD) of five floral traits and female reproductive success (under two inbreeding depression scenarios) in single-flowered hermaphroditic individuals of *P. alpina* used in the selection gradient analysis (*N* = 60). The means and SD of the subset of individuals used in the analysis of intra-floral selfing rate (*N* = 53) were almost identical; we thus present only data based on the 60 individuals.

**Table S2.** Report table showing the effects of five floral traits (standardized) on the intra-floral selfing rate of single-flowered individuals of *P. alpina* from a multivariate *glmer* model (*N* = 621 paternity-assigned seeds from 53 individuals).

Notes: n.s. *P* > 0.05, ∗ *P* < 0.05, ∗∗ *P* < 0.01, ∗∗∗ *P* < 0.001

|  | AIC | LRT | *P*-value |
| --- | --- | --- | --- |
| Full model | 240.16 |  |  |
| Stamen number | 255.52 | 17.3614 | *** |
| Pistil number | 243.94 | 5.7820 | * |
| Flowering date | 243.96 | 5.7926 | * |
| Tepal length | 238.77 | 0.6087 | n.s. |
| Stalk height | 238.55 | 0.3905 | n.s. |

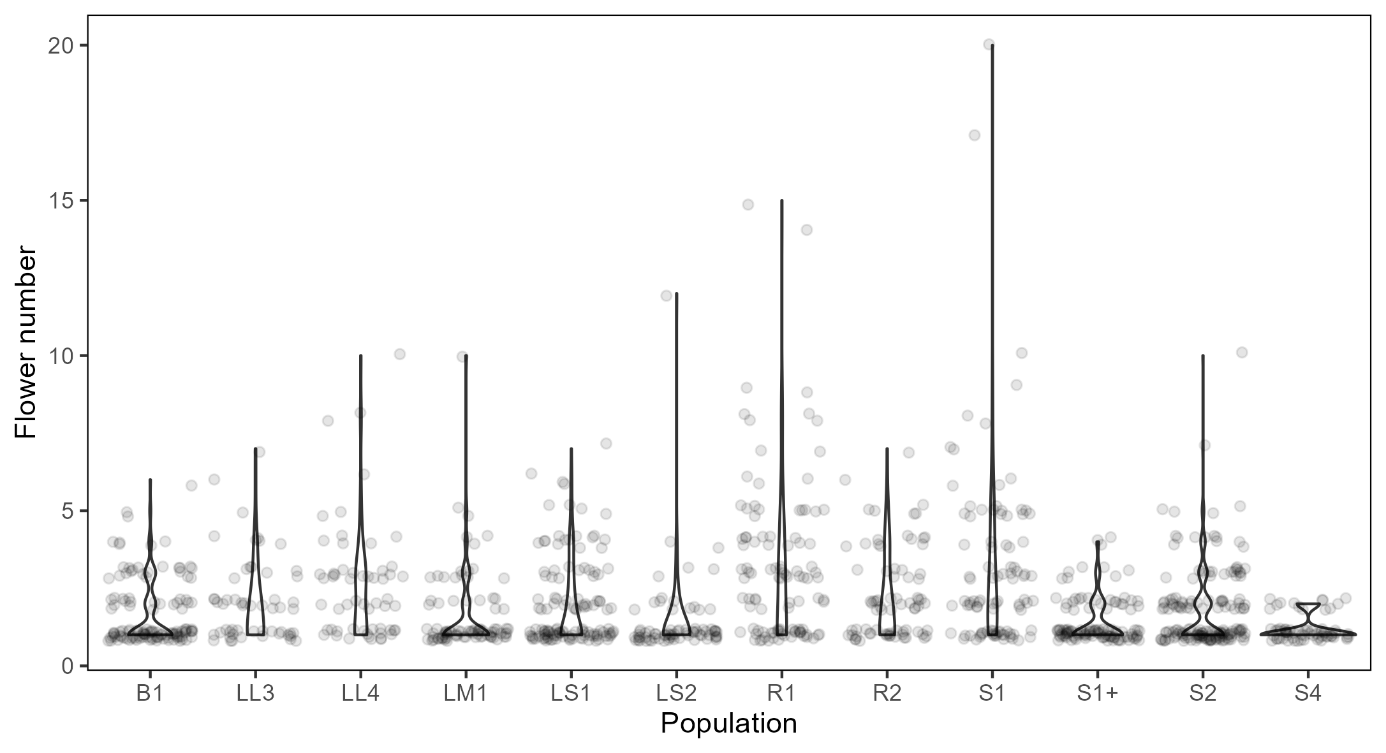

**Figure S1.** Violin plot showing the number of flowers produced by flowering individuals in 12 populations of *P. alpina* in 2022 (*N* = 1131). On average an individual produced 2.05 ± 1.76 flowers. In addition, 55% of the sampled individuals produced only a single flower. Detailed descriptions of sampled populations can be found in Chen and Pannell 2022.

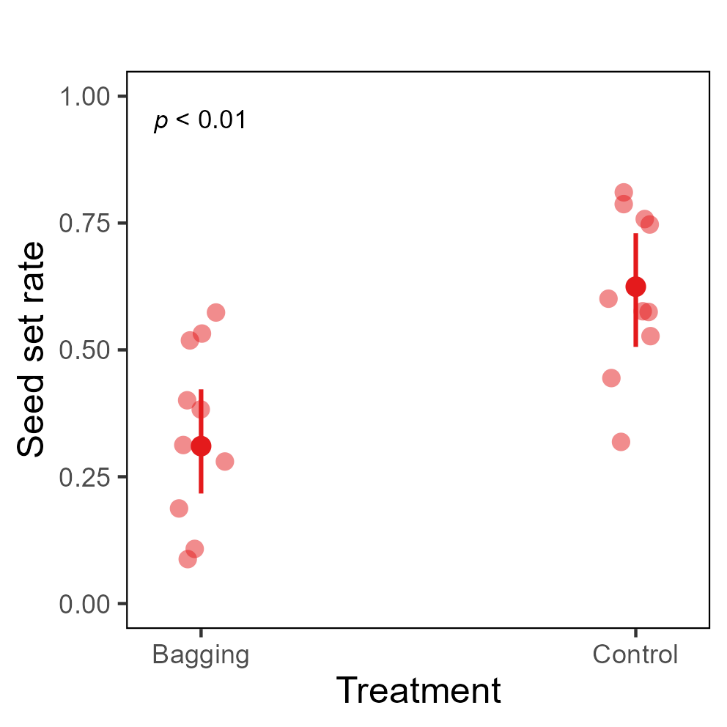

**Figure S2.** Seed set rates of bagged and intact flowers in Population LL1 in 2018 (see Chen and Pannell 2022 for details of the population). Flower buds from different individuals (*N* = 10 for each treatment) were bagged with a paper bag (Bagging treatment) or left intact (Control treatment) throughout the flowering season. The seed set rate was calculated as the number of pollinated seeds divided by the total number of pistils as described in Chen and Pannell 2022. Bagged flowers produced a significantly lower number of seeds than intact ones (*p* < 0.01), which indicates that the species is self-compatible with an incomplete dichogamy and may be capable of self-fertilizing without pollinators. However, these results should be interpreted with caution because the self-fertilization may be a consequence of touches between pistils and stamens caused by the bags under strong wind in the alpine grassland.

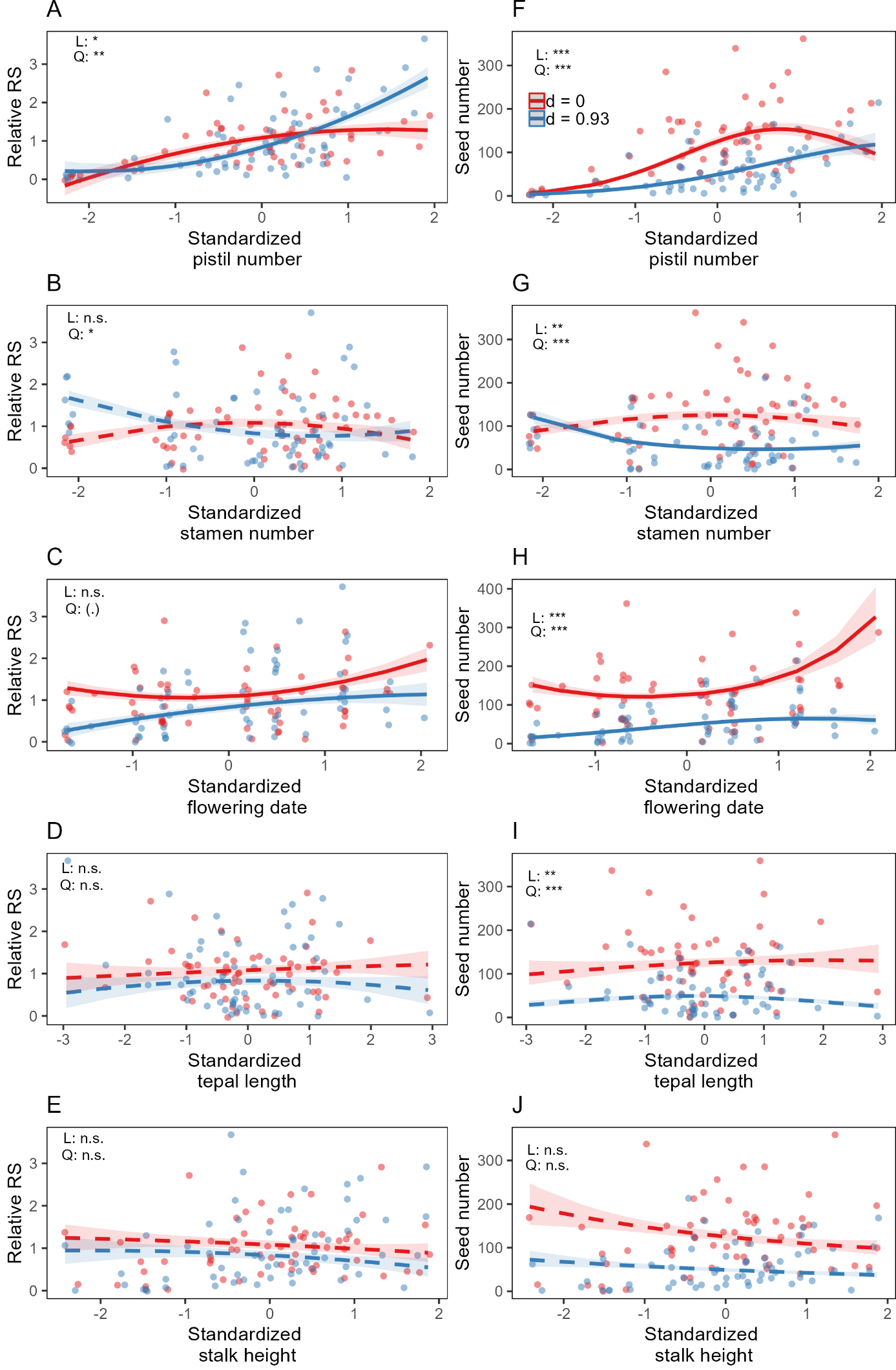

**Figure S3.** Plot showing the dependence of female fitness on pistil number (A, F), stamen number (B, G), flowering date (C, H), tepal length (D, I), and stalk height (E, J) of single-flowered hermaphroditic individuals of *P. alpina*, using relative reproductive success (RS, left-hand panel) and mature seed number (right-hand panel) as fitness proxies (*N* = 60 individuals). The two-fold transmission advantage of self-fertilization was taken into account when calculating the female fitness proxies. Each individual is represented by two points for scenarios of an inbreeding depression of zero (red) and 0.93 (blue). The shaded ribbons indicate the standard error of the regression curves. Dashed regression curves indicate non-significant linear and quadratic coefficients. The *P* values of interaction terms of inbreeding depression scenarios with linear (L) and quadratic (Q) coefficients are denoted by asteroids.

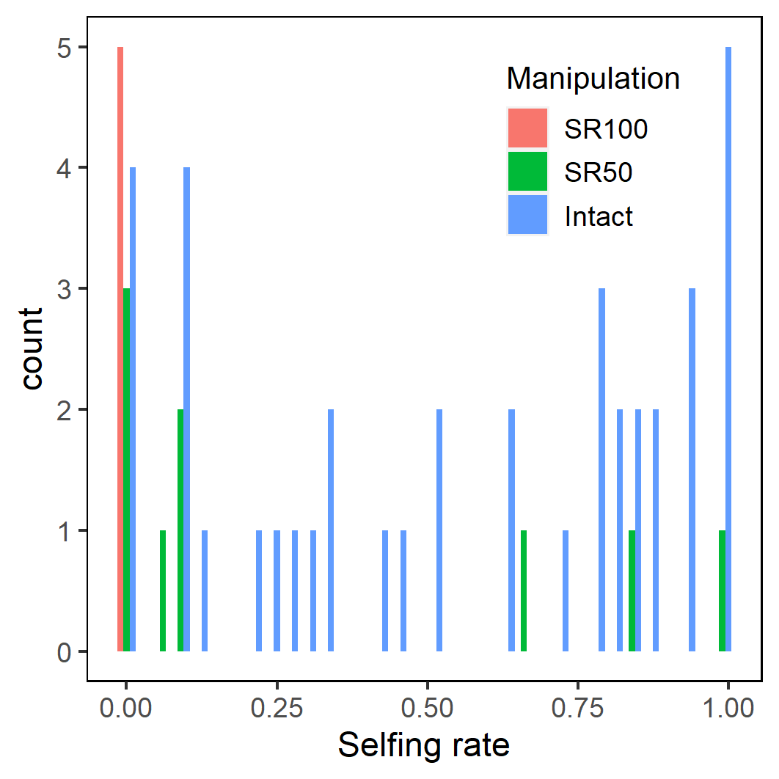

**Figure S4.** Distribution of intra-floral selfing rate of single-flowered individuals of *P. alpina* used in the analysis (*N* = 53). Individuals subject to 100 and 50 percent stamen removal manipulations are indicated by red (SR100) and green (SR50) bars, respectively.

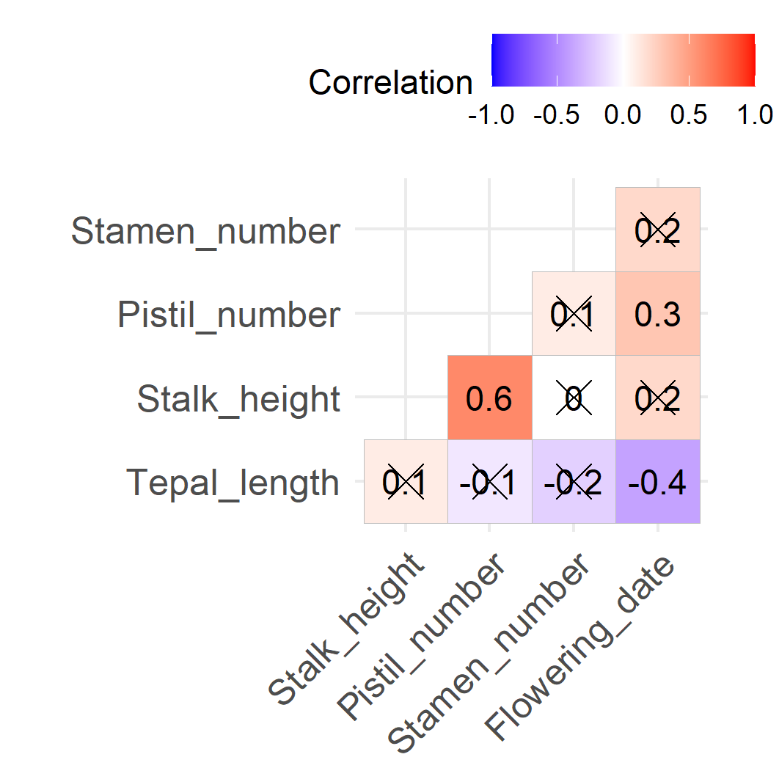

**Figure S5.** Correlation plot of the five floral traits for *P. alpina* that were subjected to selection gradient analysis (*N* = 60 hermaphroditic flowers). Non-significant (*P* > 0.05) correlations are crossed.
